# Post-outbreak surveillance strategies to support proof of freedom from foot-and-mouth disease

**DOI:** 10.1101/2021.04.27.441714

**Authors:** Richard Bradhurst, Graeme Garner, Iain East, Clare Death, Aaron Dodd, Tom Kompas

## Abstract

Whilst emergency vaccination may help contain foot-and-mouth disease in a previously FMD-free country, its use complicates post-outbreak surveillance and the recovery of FMD-free status. A structured surveillance program is required that can distinguish between vaccinated and residually infected animals, and provide statistical confidence that the virus is no longer circulating in previously infected areas.

Epidemiological models have been well-used to investigate the potential benefits of emergency vaccination during a control progam and when/where/whom to vaccinate in the face of finite supplies of vaccine and personnel. Less well studied are post-outbreak issues such as the management of vaccinated animals and the implications of having used vaccination during surveillance regimes to support proof-of-freedom. This paper presents enhancements to the Australian Animal Disease Model (AADIS) that allow comparisons of different post-outbreak surveillance sampling regimes for establishing proof-of-freedom from FMD.

A case study is provided that compares a baseline surveillance sampling regime (derived from current OIE guidelines), with an alternative less intensive sampling regime. It was found that when vaccination was not part of the control program, a reduced sampling intensity significantly reduced the number of samples collected and the cost of the post-outbreak surveillance program, without increasing the risk of missing residual infected herds.

## 1. INTRODUCTION

Emergency vaccination (as a supplement to stamping out), is increasingly being recognised as a potentially useful management strategy for foot-and-mouth (FMD) outbreaks in non-endemic countries. Vaccination may help suppress the spread of infection (Pluimers et al., 2002) and reduce the need for large-scale culling of at-risk animals (Paton et al., 2006). Epidemiological models can help inform when/where/how suppressive vaccination might best be deployed (Bates et al., 2003; Tildesley et al., 2006; Backer et al., 2012a; Tildesley et al., 2012; Boklund et al., 2013; Porphyre et al., 2013; Hayama et al., 2013; Durr et al., 2014; Rawdon et al., 2018; Marschik et al., 2020), and how constraints on vaccine supply and/or response personnel might impact the effectiveness of control (Abdalla et al., 2005; Roche et al., 2014).

The introduction of vaccination in a previously FMD-free jurisdiction, does, however, complicate post-outbreak surveillance and the recovery of FMD-free status. This is an important issue for countries that have significant exports of livestock and livestock products. To apply for FMD-free status after eradicating an FMD outbreak, a country has to present a case, in the form of a dossier, to the World Organization for animal Health (OIE) (USDA, 2015; OIE, 2016). The dossier should include extensive descriptions of the livestock populations and management systems, animal health services (both field and laboratory), and detailed information on how the outbreak was managed (including the relevant legislation and a chronology of outbreak events and implemented control measures). A key element underpinning the case for freedom from FMD is a structured post-outbreak surveillance program that provides statistical confidence that the virus is no longer circulating in previously infected areas.

Animals infected with FMD virus (FMDV) produce antibodies to both viral structural proteins and non-structural proteins. Vaccinated animals produce antibodies mainly or entirely to the viral structural proteins. Structural protein (SP) serological tests can thus be used to screen unvaccinated populations for evidence of FMDV exposure or infection. These tests are serotype-specific and are highly sensitive provided that the virus or antigen used in the test is closely matched to the strain circulating in the field (Constable et al., 2017). If vaccinated animals are retained in the post-outbreak population then surveillance to support regaining FMD-free status will be much more difficult. Vaccinated animals will test positive using standard structural protein (SP) serological tests (OIE, 2012; OIE, 2016), and some that have been exposed to infection, especially soon after vaccination, may become infected then recover, or go on to become carriers. Vaccinated animals that are subsequently infected can be identified using FMD non-structural protein (NSP) tests (OIE, 2016). It is important to identify NSP-positive animals in a vaccinated population, even if they are not carriers, as they may cause problems for market access in the future. As vaccinated animals exposed to infection may become sub-clinically and persistently infected it is necessary to find and remove all infected vaccinated animals in order to regain FMD–free status (Paton et al., 2006; Paton et al., 2014).

The purpose of post-outbreak surveillance is to detect acute or persistent infection, especially sub-clinical infections, and recovered animals that may have been missed during the control program. In this paper, herds containing recovered animals or carriers at the end of the outbreak are referred to as ‘residual herds’. Assessing the disease-free status of a country may be achieved through random survey and/or non-survey-based surveillance, using classical statistical or Bayesian methods (Caporale et al., 2012). Survey-based approaches have generally been preferred because they are considered more objective and defensible and a quantitative estimate of confidence of a disease being absent from a population can be readily calculated (Caporale et al., 2012).

Ideally, post-outbreak surveillance should locate residual herds without being unnecessarily complex, expensive, and time consuming. All herds that test positive (i.e., both true positives and false positives), will generate workload to clarify their actual status. Due to the major trade implications associated with falsely declaring freedom from infection, any positive result is followed up with a regime of further testing. Under the OIE Code (OIE, 2016), finding evidence of infection in the target population automatically invalidates any claim for freedom.

Epidemiological modelling tools in a non-endemic setting tend to focus on the period from the introduction of infection through to the completion of the control program. As such, the post-outbreak consequences of having used vaccination during a control program are less well studied. From a policy perspective it would be very useful if disease managers had access to decision support tools that could be used to evaluate surveillance strategies for the recovery of FMD-free status. The Australian Animal Disease spread model (AADIS) (Bradhurst et al., 2015) is a national-scale epidemiological model used by animal health authorities in Australia to support FMD planning and preparedness. AADIS has previously been upgraded to include post-outbreak management strategies for vaccinated animals (retention, waste, salvage) and the consequences on recovery of FMD-sensitive markets per OIE guidelines (Bradhurst et al., 2019). In this paper we present model enhancements to assist with the evaluation of post-outbreak surveillance approaches and show, using a case study, how the model can be used to compare different approaches to surveillance in terms of effectiveness, resource requirements and costs

## 2. MATERIALS AND METHODS

### 2.1. The AADIS epidemiological model

The AADIS model is a spatiotemporal agent-based simulation of the spread and control of emergency animal disease (Bradhurst et al., 2015). Each herd agent has an embedded set of differential equations that model the herd’s infected, infectious, serological and clinical prevalence over time, taking into account species, production system and virus strain. For this study we have assumed a Type O Pan-Asia strain. The agents interact in a model environment that stochastically spreads disease across multiple spread pathways (direct contacts, indirect contacts, saleyard spread, airborne transmission and local spread). Control measures (stamping out, surveillance, tracing, movement restrictions and vaccination), are implemented per the Australian Veterinary Emergency Plan (AUSVETPLAN) for FMD (Animal Health Australia, 2014). As in an actual outbreak response, the AADIS control measures are dynamically constrained by the availability of resources (responders and consumables such as vaccine), the accuracy of reports of clinical disease (as false reports still consume surveillance resources), inefficiencies in tracing systems, and non-conformance to movement restrictions. The AADIS model is able to efficiently represent national-scale epidemics due to the hybrid model architecture, concurrent software architecture, in-memory database, and grid-based spatial indexing (Bradhurst et al., 2016).

### 2.2. Incorporating post-outbreak surveillance

A new AADIS post-outbreak management module was developed to simulate serological surveillance to support proof-of-freedom in terms of:

a. a sampling regime for selecting number herds to test, and the number of samples to take from each test herd in a target population
b. a testing regime that defines which serological test (or combination of tests) to employ, taking into account test sensitivity (S_e_) and specificity (S_p_)

This module allows surveillance strategies to be compared in terms of:

- effectiveness in finding residual herds
- number of false positive reactors generated
- time taken to complete the surveillance program
- resources required and overall cost (field teams, laboratory tests, reagents)

AADIS post-outbreak surveillance is defined in terms of ‘clusters’ of discrete infected areas (Anon., 2007), where a cluster is defined as the set of premises enclosed by one or more intersecting controlled areas. Under the Australian Veterinary Emergency Plan AUSVETPLAN (Animal Health Australia, 2014), two types of controlled areas are used:

1. The Restricted Area (RA) defined by a radial buffer (minimum 3 km) around an infected premises (IP). The RA imposes the highest levels of livestock movement restrictions.
2. The Control Area (CA) defined by an annulus with inner radius (default 3 km) and outer radius (minimum 10 km) around an IP (i.e., enclosing the RA). The CA has lower levels of movement restrictions than the RA.

An example is provided in Figure 1 where five clusters have been defined after a hypothetical FMD outbreak. Each cluster is comprised of a set of one or more IPs (black dots), the premises in the aggregated RA (shaded red) enclosing the IPs, and the premises in the aggregated CA (shaded green). Surveillance sampling regimes are applied to each cluster independently.

**Figure 1.**
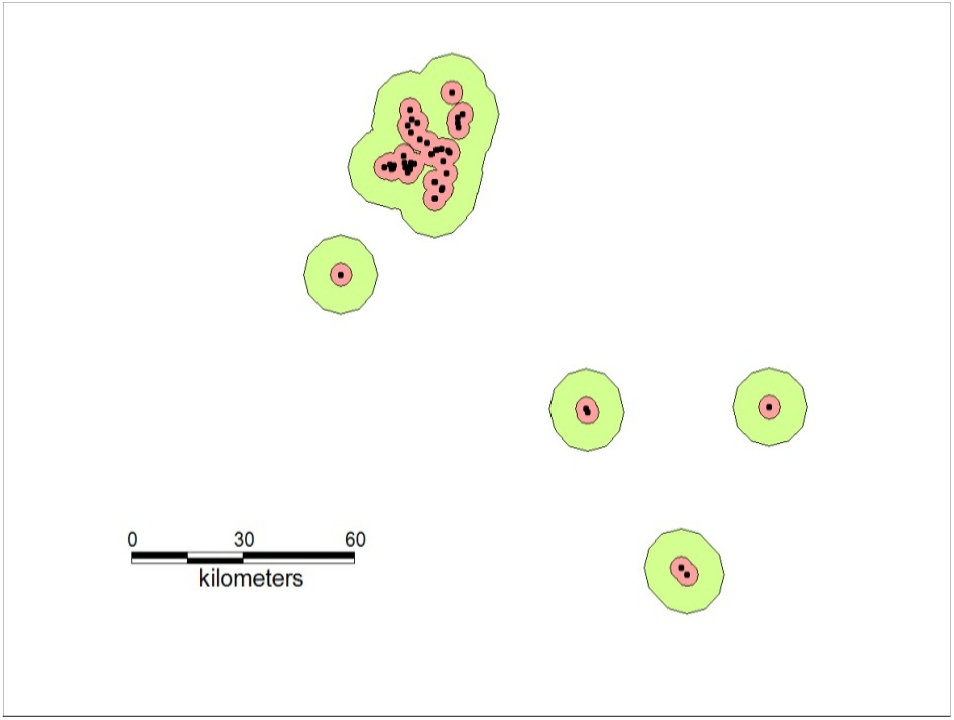
Post-outbreak clusters (n=5) associated with a hypothetical FMD outbreak.

#### 2.2.1. Sampling regime

It is not possible to prove that infection (past or present) is not in a population. Rather, freedom from infection is probabilistic with a level of confidence that disease is not present at a specified minimum level, known as the ‘design prevalence’. There are several techniques for determining the minimum sample size necessary to detect the presence of infection (Canon and Roe, 1982; Cameron and Baldock, 1998; Sergeant, 2017). AADIS employs Cannon and Roe’s (1982) binomial approximation for calculating sample size.

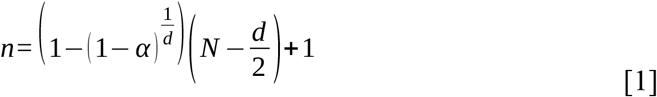

where:

*n* = sample size

*N* = herd size

*d* = number of target positives in herd (= design prevalence * test S_e_ * N)

*α* = confidence level

if *n* > *N* then *n* = *N*

A sampling regime is described in terms of ‘confidence level:design prevalence’, for example, a within-herd sampling regime of 95:5 involves testing sufficient animals (per equation 1) to achieve 95% confidence that an infected prevalence of at least 5% would be detected. Note that equation 1 can also be used to calculate the number of herds to sample within a target population of herds in order to achieve a design prevalence with a specified confidence.

The AADIS model allows the user to set sampling regimes for different species (cattle, sheep and pigs) and for previous RAs and CAs. In the absence of an international standard, the baseline sampling regime (Table 1) is based on the European approach (European Union, 2003; Anon., 2007). Within each RA cluster, all sheep flocks are tested by serology with a 95:5 within-flock sampling regime. Surveillance of cattle and pigs in an RA cluster is based on clinical inspections of all animals in all herds, i.e., no specific sample collection and serological testing is required. However, while this may be adequate for detecting evidence of active infection, it provides little assurance that recovered cattle herds, potentially containing carrier animals, have not been missed during the control program. Accordingly, the option of serological sampling and testing cattle according to a user specified within-herd sampling regime is included (default: 95:5).

**Table 1.** Diagnostic test parameter settings.

| Test | Type | Species | Control program with vaccination | | Control program without vaccination | | Cost (A\$)<sup>4</sup> |
| --- | --- | --- | --- | --- | --- | --- | --- |
|  |  |  | Sensitivity | Specificity | Sensitivity | Specificity |  |
| C-ELISA | SP<sup>1</sup> | cattle | - | - | 0.99 | 0.99 | 35 |
| C-ELISA | SP<sup>1</sup> | sheep | - | - | 0.99 | 0.99 | 35 |
| C-ELISA | SP<sup>1</sup> | pigs | - | - | 0.99 | 0.82 | 35 |
| C-ELISA | SP<sup>1</sup> | other<sup>3</sup> | - | - | 0.99 | 0.94 | 35 |
| 3ABC-ELISA | NSP<sup>2</sup> | cattle | 0.8 | 0.99 | 0.93 | 0.99 | 35 |
| 3ABC-ELISA | NSP<sup>2</sup> | sheep | 0.8 | 0.99 | 0.9 | 0.99 | 35 |
| 3ABC-ELISA | NSP<sup>2</sup> | pigs | 0.7 | 0.99 | 0.73 | 0.99 | 35 |
| 3ABC-ELISA | NSP<sup>2</sup> | other<sup>3</sup> | 0.77 | 0.99 | 0.85 | 0.99 | 35 |

<sup>1</sup>structural protein antibody test
<sup>2</sup>non-structural protein antibody test
<sup>3</sup>any species present in a smallholder herd (defined as 50 animals no less than 20 hectares)
<sup>4</sup>Australian dollars

Within each CA cluster, sufficient sheep flocks are randomly selected to meet a 95:2 sampling regime. Within each selected flock, sufficient animals are tested to meet a 95:5 sampling regime. Surveillance of cattle and pigs in each RA cluster is based on clinical inspections of all animals in all herds. There is also the option of including 95:2 herd sampling and 95:5 within-herd sampling.

Additional surveillance is required when vaccination is used during the control program and vaccinated animals are not removed from the population. Under the European approach, all vaccinated herds are tested excluding those that went on to become IPs. Surveillance involves clinical inspection of all susceptible animals in all herds, and NSP testing of vaccinated cattle and sheep herds, which are tested according to a 95:5 within-herd sampling regime.

As with other disease control processes in AADIS, post-outbreak surveillance is subject to resource constraints. Potential limitations are the sample collection rate (dependent on the number of surveillance teams and how long it takes to sample each herd), and the sample test rate (dependent on laboratory throughput).

#### 2.2.2. Testing regime

It is unusual to base surveillance on a single test result. In practice, any sample that tests positive in an initial (screening) test would be subject to a second confirmatory test (OIE 2016; Brocchi et al., 2006; Paton et al., 2006). A serial screening and confirmatory test process increases specificity but reduces overall sensitivity of the testing process which needs to be taken into account. AADIS allows the user to define test pairs [screening, confirmatory tests]. In this study, the test pair for an unvaccinated target population is [C-ELISA, 3ABC-ELISA], and for a vaccinated population is [3ABC-ELISA, 3ABC-ELISA]. Model parameter values for the serological tests used in this study are provided in Table 1. This information was provided by staff from the Australian Animal Health Laboratory (J Watson pers. comm. May 2017).

### Test results

#### (a) True positive

The probability of infection being detected in a residual herd (i.e., a true positive test result), depends on the within-herd seroprevalence, the test sensitivity, and the number of animals sampled (Canon and Roe, 1982).

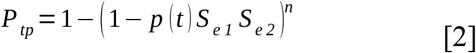

where

*P*_*tp*_ = probability of a true positive test result (i.e., at least one animal tests positive)

*p(t)* = herd seroprevalence on the testing day *t*

*S*_*e1*_ = sensitivity of the screening test

*S*_*e2*_ = sensitivity of the confirmatory test

*n* = number of animals sampled in the herd

For simplicity, equation 2 assumes that the test results are independent and does not take sensitivity co-variance (the probability of both tests being positive), into account (Paton et al., 2006; Brocchi et al., 2006).

#### (b) False positive

The probability of infection being falsely detected in a non/never-infected herd (i.e., a false positive test result), depends on the test specificity (Canon and Roe, 1982).

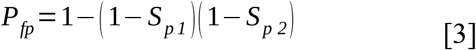

where

*P*_*fp*_ = probability of a false positive result (i.e., at least one animal tests positive)

*S*_*p1*_ = specificity of the screening test

*S*_*p2*_ = specificity of the confirmatory test

For simplicity, equation 3 assumes that the test results are independent and does not take specificity co-variance (the probability of both tests being negative), into account (Paton et al., 2006; Brocchi et al., 2006).

#### (c) True negative

As AADIS is a simulation that explicitly models within-herd prevalence (infected, serological and clinical), the number of true negative test results is calculated as the number of actual non/never-infected herds minus the number of false positives.

#### (d) False negative

The number of false negative test results is the number of residual herds minus the number of true positives.

### 2.3. Cost estimates and assumptions

Costs are incurred through:

- Coordination of surveillance activities through the continued operation of Local and State Disease Control Centres beyond the control phase. The cost of operating a disease control centre is estimated at A$120,000 (Australian dollars) per centre per day, and takes into account expenses such as salaries including penalty rates, fixed overheads such as rent, operating expenses such as meals, and accommodation (Kevin Cooper, pers. comm., 2012; Steven Riley, pers. comm., 2017).
- Clinical inspections and collection of samples - based on the time and materials required to muster and inspect an ‘average’ herd of each type. A surveillance team is assumed to comprise two people (one professional, plus one assistant), who inspect on average 500 animals per day. The cost of a surveillance visit (excluding laboratory tests) for each herd type is shown in Table 2. They assume a daily salary cost of A$1500 plus a fixed cost of A$100 for travel and A$150 for disposables (per herd).
- Laboratory tests - calculated by multiplying the number of samples collected from each herd (dependent on the sampling strategy), by the cost of the tests used (Table 1).
- Following-up positive test results. This is not currently included in the AADIS cost reporting.

**Table 2.**
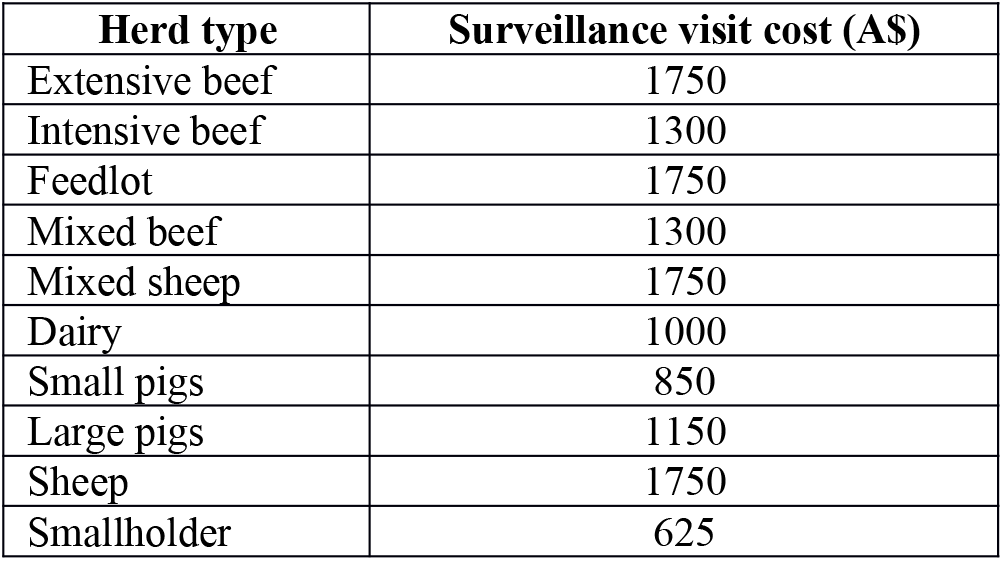
Cost of a surveillance visit (excluding laboratory costs) for each herd type.

### 2.4. Case study

To demonstrate the functionality, a case study is presented in which the baseline post-outbreak sampling regime based on the EU approach is compared with a hypothetical reduced sampling regime (Table 3).

**Table 3.**
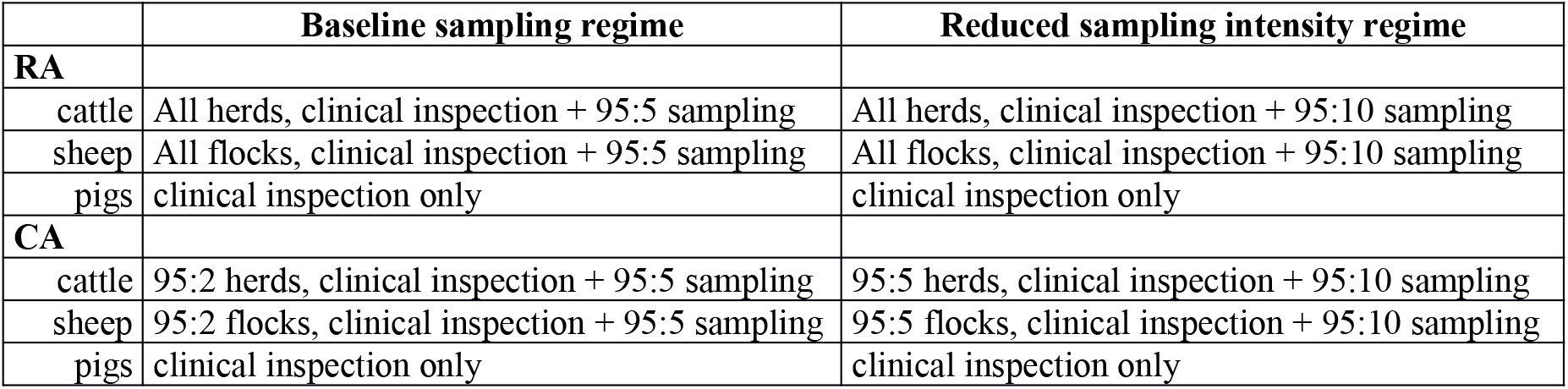
Post-outbreak sampling regimes under baseline and reduced sampling intensity approaches

|  | Baseline sampling regime | Reduced sampling intensity regime |
| --- | --- | --- |
| **RA** |  |  |
| cattle | All herds, clinical inspection + 95:5 sampling | All herds, clinical inspection + 95:10 sampling |
| sheep | All flocks, clinical inspection + 95:5 sampling | All flocks, clinical inspection + 95:10 sampling |
| pigs | clinical inspection only | clinical inspection only |
| **CA** |  |  |
| cattle | 95:2 herds, clinical inspection + 95:5 sampling | 95:5 herds, clinical inspection + 95:10 sampling |
| sheep | 95:2 flocks, clinical inspection + 95:5 sampling | 95:5 flocks, clinical inspection + 95:10 sampling |
| pigs | clinical inspection only | clinical inspection only |

FMD virus of type O is assumed to be introduced in a piggery through illegal feeding of swill containing infectious material sourced from overseas. The outbreak begins, in September, on a small 25-sow pig farm (n=247 pigs) just west of Leongatha in Gippsland, Victoria (38.4740°S, 145.9437°E). FMD is recognised and reported to the authorities 14 S, 145.9437°S, 145.9437°E). FMD is recognised and reported to the authorities 14 E). FMD is recognised and reported to the authorities 14 days after introduction. AADIS was run (1000 iterations) for the ‘silent spread’ phase A single iteration, consistent with the median number of infections from the 1000 runs, was selected (Figure 2). There are 18 infected herds in the population. This iteration serves as a common starting point for the control programs under test (from day 15 onward), for all subsequent runs in the case study.

**Figure 2.**
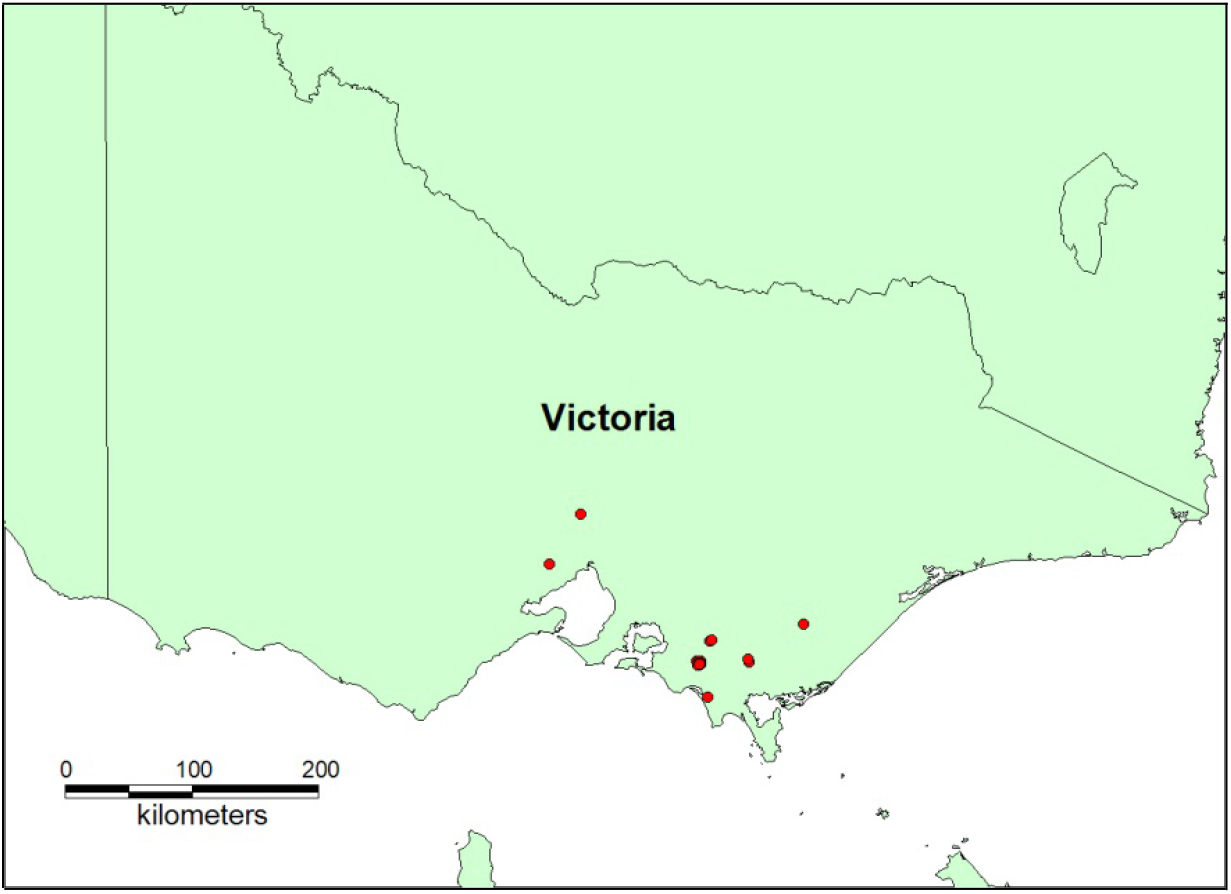
Infected herds (n=18) in the VIC outbreak scenario at the end of the silent spread phase

Control programs with and without vaccination were simulated. Selected configuration parameter settings for the case study are provided in Table 4.

**Table 4.**
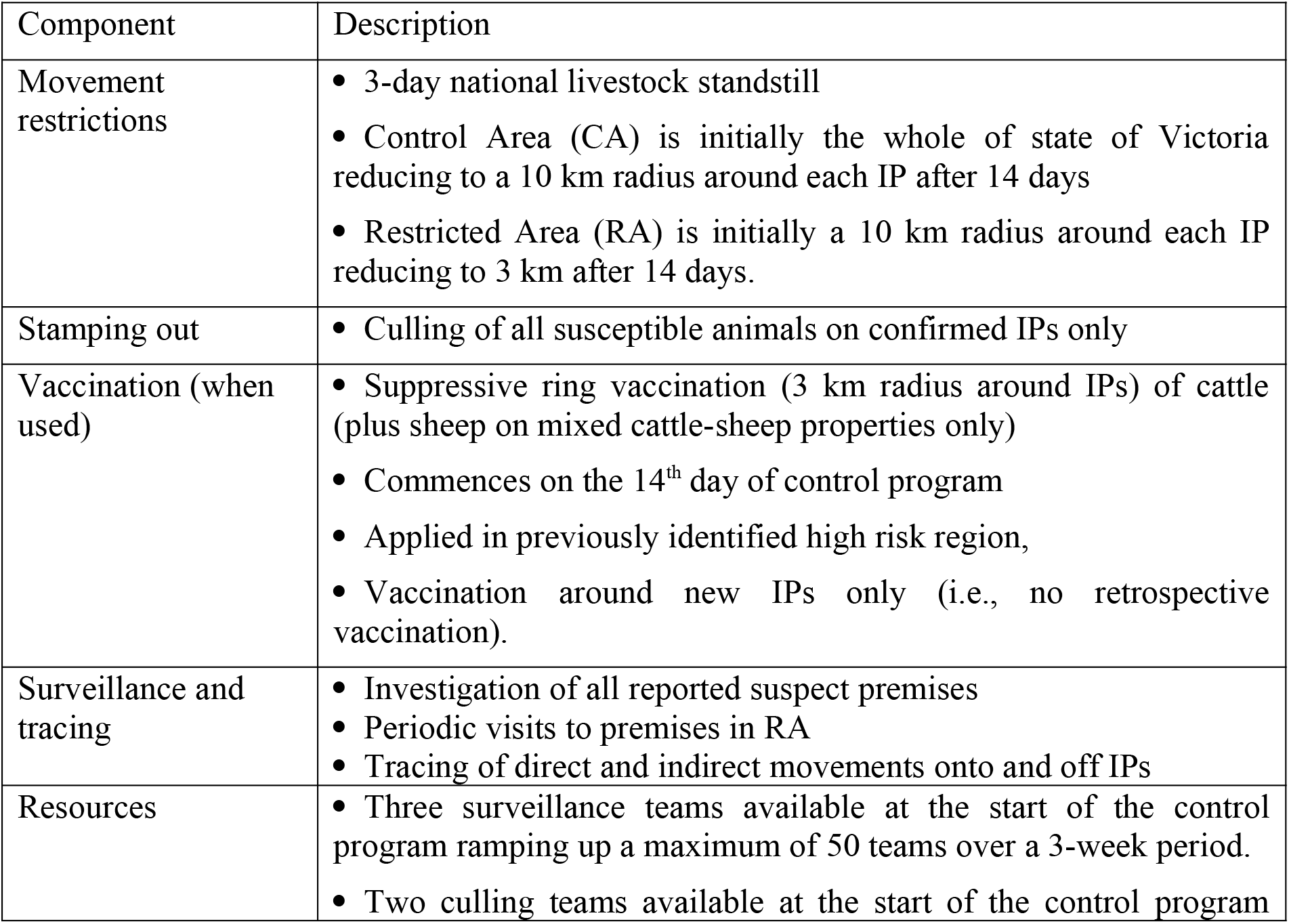
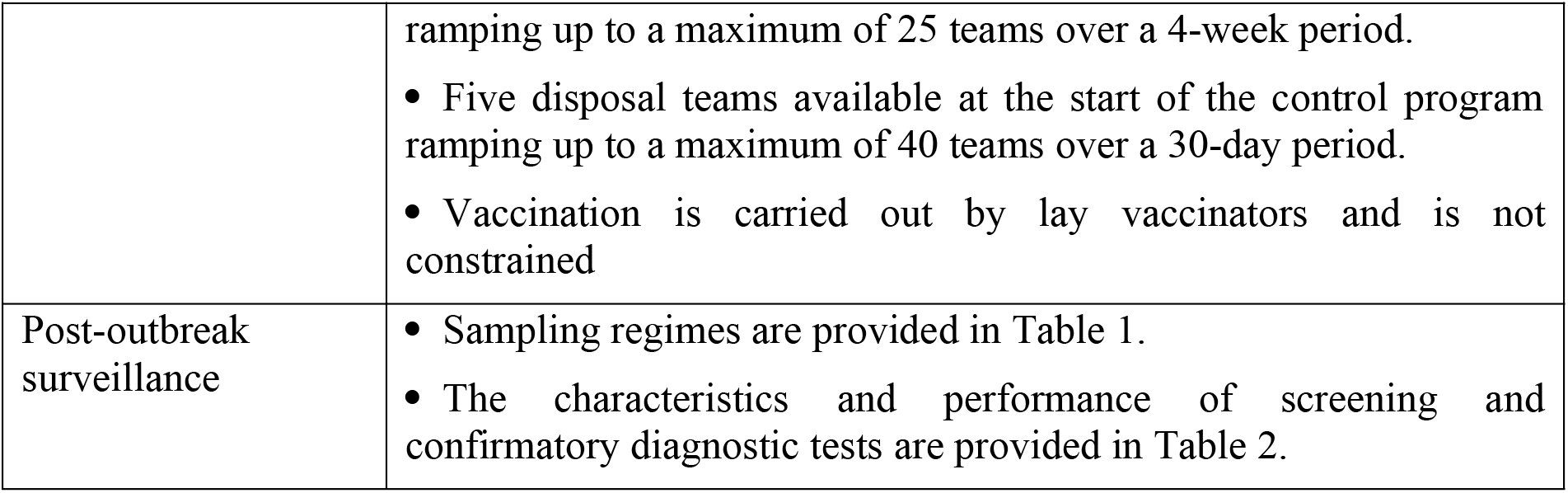
Control parameter settings for the case study

| Component | Description |
| --- | --- |
| Movement restrictions | • 3-day national livestock standstill<br>• Control Area (CA) is initially the whole of state of Victoria reducing to a 10 km radius around each IP after 14 days<br>• Restricted Area (RA) is initially a 10 km radius around each IP reducing to 3 km after 14 days. |
| Stamping out | • Culling of all susceptible animals on confirmed IPs only |
| Vaccination (when used) | • Suppressive ring vaccination (3 km radius around IPs) of cattle (plus sheep on mixed cattle-sheep properties only)<br>• Commences on the 14<sup>th</sup> day of control program<br>• Applied in previously identified high risk region,<br>• Vaccination around new IPs only (i.e., no retrospective vaccination). |
| Surveillance and tracing | • Investigation of all reported suspect premises<br>• Periodic visits to premises in RA<br>• Tracing of direct and indirect movements onto and off IPs |
| Resources | • Three surveillance teams available at the start of the control program ramping up a maximum of 50 teams over a 3-week period.<br>• Two culling teams available at the start of the control program |
|  | ramping up to a maximum of 25 teams over a 4-week period.<br>• Five disposal teams available at the start of the control program ramping up to a maximum of 40 teams over a 30-day period.<br>• Vaccination is carried out by lay vaccinators and is not constrained |
| Post-outbreak surveillance | • Sampling regimes are provided in Table 1.<br>• The characteristics and performance of screening and confirmatory diagnostic tests are provided in Table 2. |

### 2.5. Study design and statistical methods

One thousand iterations of the case study outbreak scenario were run for each of the following control and post-outbreak surveillance combinations:

i. stamping out only control with the (EU-based) baseline post-outbreak surveillance sampling regime
ii. stamping out only control with the alternate (less intensive) post-outbreak surveillance sampling regime
iii. stamping out plus suppressive ring vaccination control with the (EU-based) baseline post-outbreak surveillance sampling regime
iv. stamping out plus suppressive ring vaccination control with the alternate (less intensive) post-outbreak surveillance sampling regime

Simulations were run until the control and post-outbreak management program was complete, or for 365 days, whichever came first.

The baseline and alternate (reduced intensity) sampling regimes were compared, for both the vaccination and non-vaccination control programs, using the following model outputs:

- days taken to complete the surveillance program
- number of herds tested
- number of samples taken
- number of clinical inspections carried out
- cost of the surveillance program (A$)
- number of true/false positive test results (i.e., herds requiring follow-up)
- number of false negative test results (i.e., residual herds that were not detected)

The DPlot add-on (Hydesoft Computing, Vicksburg, MS, USA) for Excel (Microsoft Corp. USA) was used to produce box and whisker plots. The box represents the 25 – 75 percentiles range. The horizontal band within the box represents the median. The whiskers represent the 0 – 25 percentile (lower) and the 75 – 100 percentile (upper).

Where comparisons between different approaches were made, data were analysed using STATA statistical software package (StataCorp, 2017). Initially all data was examined for normality using: (i) visual appraisal of histograms of the data, (ii) determination of the skew and kurtosis of each data set and its deviation from the values expected in a normal distribution, and (iii) an automated search of a subset of the ladder of powers for a transform that converted the data to normality. All of the data sets were non-normal. No standard transformations transformed the data into a normal distribution. All data sets were log transformed to minimise the over-distribution (left skew) observed.

Data sets were compared using both the one-way ANOVA and the Kruskal-Wallis tests. The one-way ANOVA has been reported as robust to deviations from normality when the data sets are large and was used because it is a more rigorous test (Feir-Walsh and Toothaker, 1974; Schmider et al., 2010). In addition, pairwise comparisons of datasets could be done automatically when more than two groups were being compared. To examine the impact of the data being non-normal, all results were checked using the Kruskal-Wallis test. The one-way ANOVA and the Kruskal-Wallis test produced comparable results for each comparison examined.

## 3. RESULTS

### 3.1. Control program without vaccination

Results of the comparison between the baseline and alternate (reduced intensity) surveillance sampling regimes after a stamping out only control program are provided in Figure 3 and Table 5. Under the assumptions used for this study, the post-outbreak surveillance program took around 6.5 weeks to complete for both sampling regimes. As all herds in previously infected areas still required clinical inspection, sampling intensity had no significant effect on the time required to complete the surveillance program.

**Table 5.** Summary of baseline and alternate sampling regimes after non-vaccination control program.

| Model outcome variable | Baseline sampling regime<br>(mean ± stdev) | Reduced intensity sampling regime<br>(mean ± stdev) |
| --- | --- | --- |
| Surveillance duration (days) | 46.2 ± 13.1<sup>a†</sup> | 44.5 ± 14.0<sup>a</sup> |
| Herds tested | 1811 ± 505<sup>a</sup> | 1698 ± 527<sup>b</sup> |
| Samples collected | 87719 ± 23931<sup>a</sup> | 45042 ± 13819<sup>b</sup> |
| Clinical inspections | 3200 ± 983<sup>a</sup> | 3133 ± 1053<sup>a</sup> |
| Surveillance cost (A\$ million) | 6.35 ± 1.81<sup>a</sup> | 4.79 ± 1.54<sup>b</sup> |
| Herds that tested positive | 8.77 ± 3.66<sup>a</sup> | 4.57 ± 2.61<sup>b</sup> |
| Residual herds missed | 0<sup>a</sup> | 0<sup>a</sup> |

<sup>†</sup> Within rows, figures with differing superscripts are significantly different ( $p < 0.05$ )

**Figure 3.**
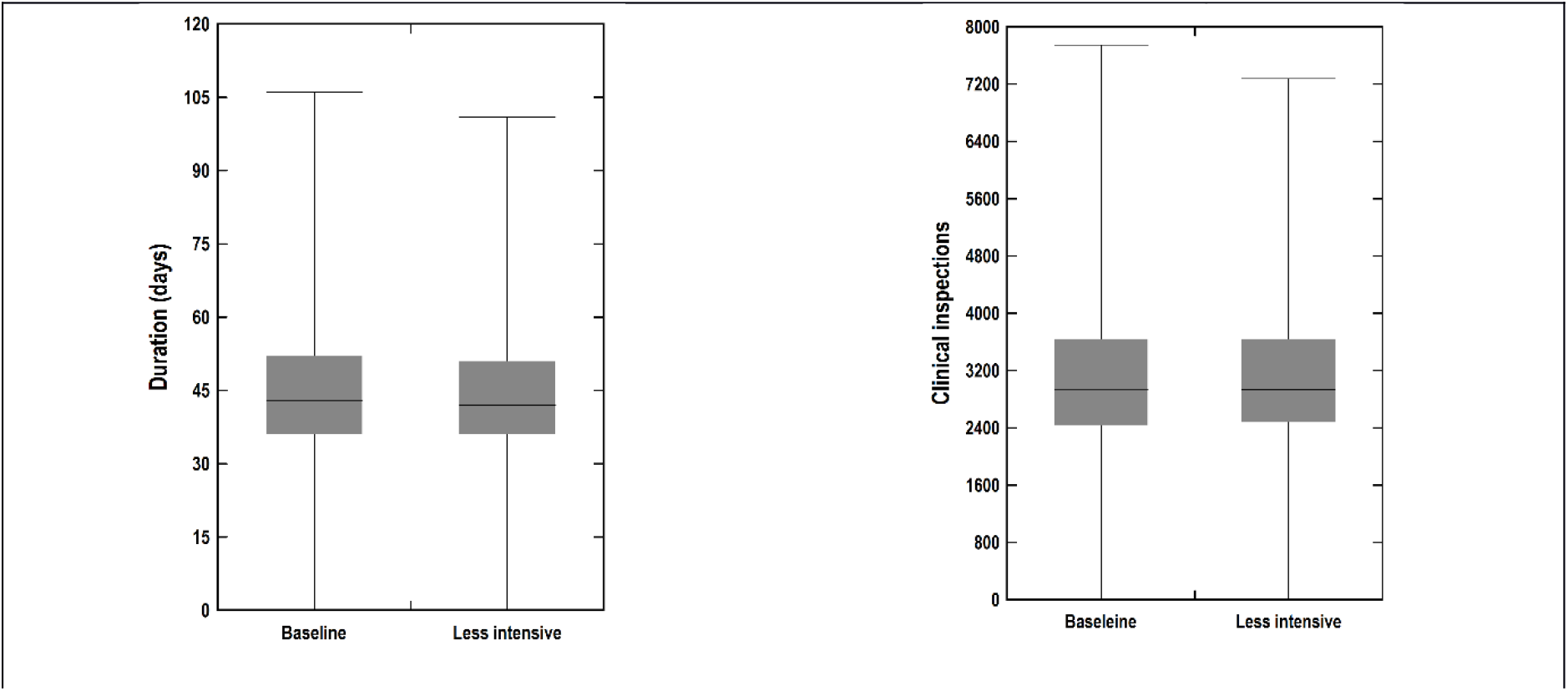
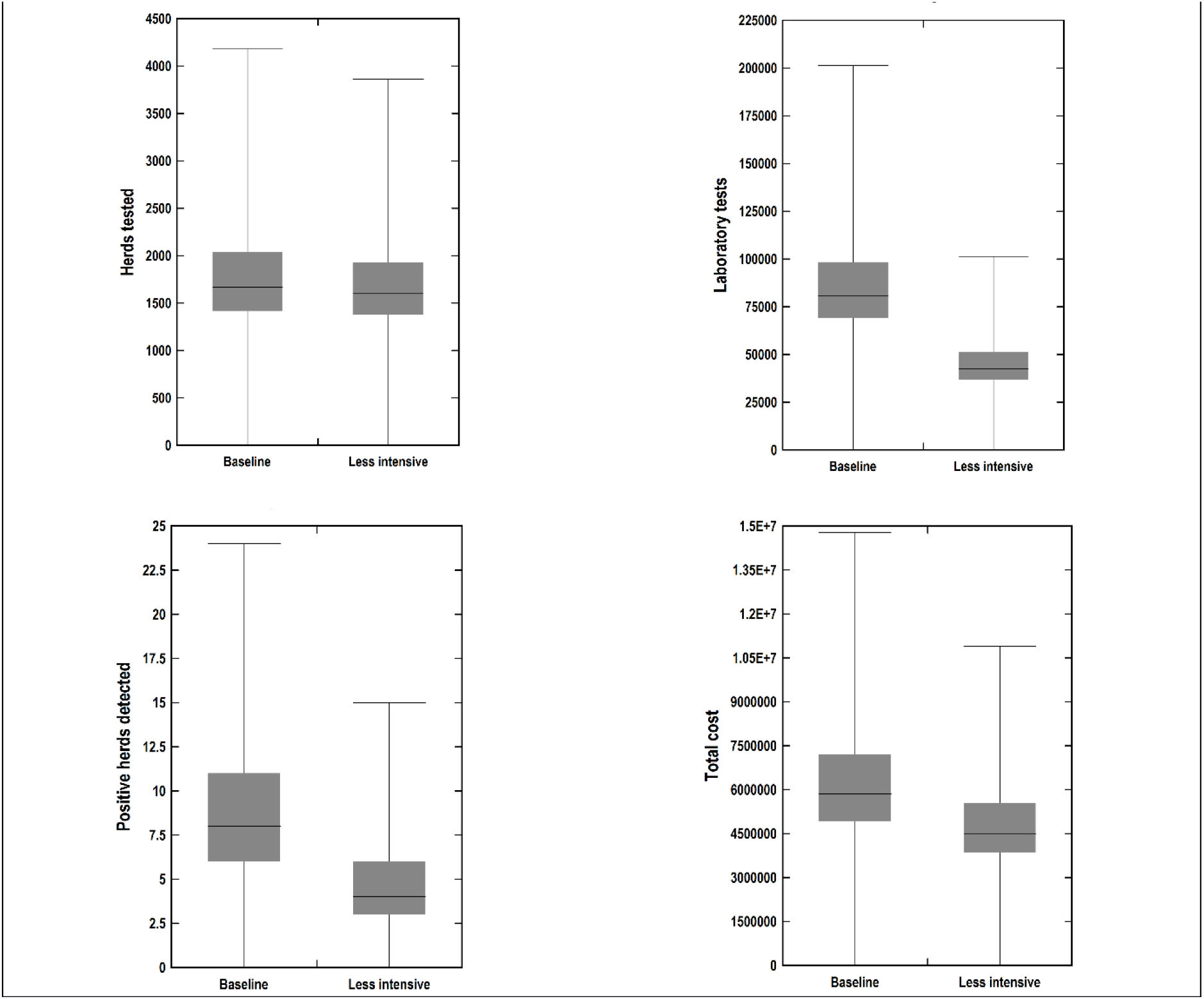
Comparison of baseline and alternate sampling regimes after non-vaccination control program

The reduced intensity sampling regime significantly reduced the number of herds tested, the number of samples collected, the cost of post-outbreak surveillance, and the number of positive herds requiring follow-up (on average by 6%, 49%, 25% and 48% respectively), compared to the baseline approach. There were no residual infected herds in the population and as such the reduction in sampling intensity could have no effect the number of false negative test results.

### 3.2. Control program with vaccination

Results of the comparison between the baseline and alternate (reduced intensity) surveillance sampling regimes after a control program that includes vaccination are provided in Figure 4 and Table 6. Under the assumptions used for this study, the post-outbreak surveillance program took around 6 weeks to complete for both sampling regimes. As all herds in previously infected areas still required clinical inspection, sampling intensity had no significant effect on the time required to complete the surveillance program.

**Table 6.** Summary of baseline and alternate sampling regimes after vaccination control program.

| Model outcome variable | Baseline sampling<br>(mean ± stdev) | Reduced intensity sampling<br>(mean ± stdev) |
| --- | --- | --- |
| Surveillance duration (days) | 43.2 ± 10.4<sup>a†</sup> | 42.2 ± 10.4<sup>b</sup> |
| Herds tested | 1709 ± 411<sup>a</sup> | 1593 ± 378<sup>b</sup> |
| Samples collected | 82879 ± 19502<sup>a</sup> | 42300 ± 9900<sup>b</sup> |
| Clinical inspections | 2954 ± 752<sup>a</sup> | 2971 ± 803<sup>a</sup> |
| Surveillance Cost (A\$ million) | 5.93 ± 1.44<sup>a</sup> | 4.53 ± 1.15<sup>b</sup> |
| Herds that tested positive | 9.22 ± 3.92<sup>a</sup> | 4.6 ± 2.50<sup>b</sup> |
| Residual herds missed | 1.86 ± 2.01<sup>a</sup> | 1.40 ± 1.53<sup>a</sup> |

<sup>†</sup> Within rows, figures with differing superscripts are significantly different ( $p < 0.05$ )

**Figure 4.**
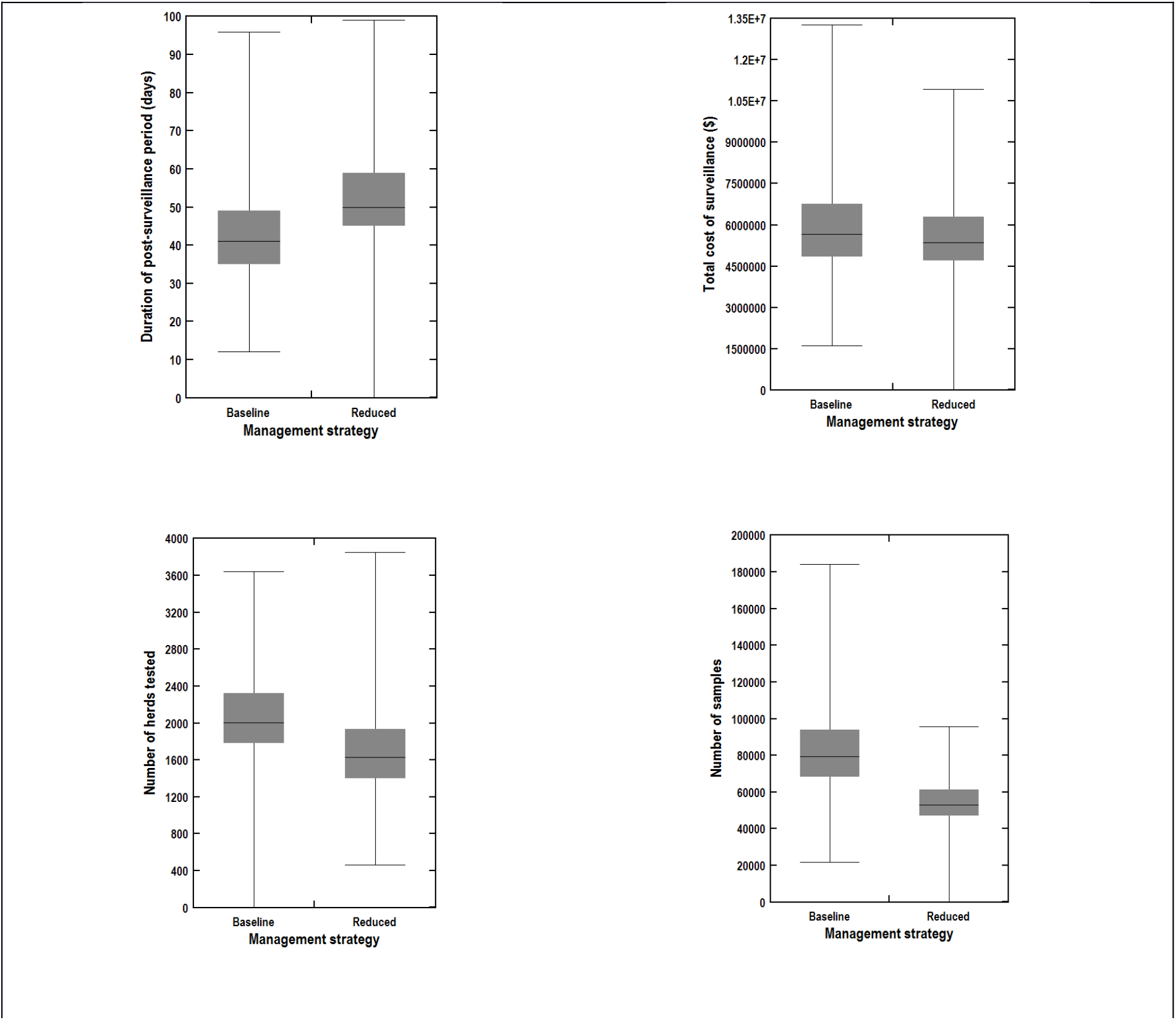
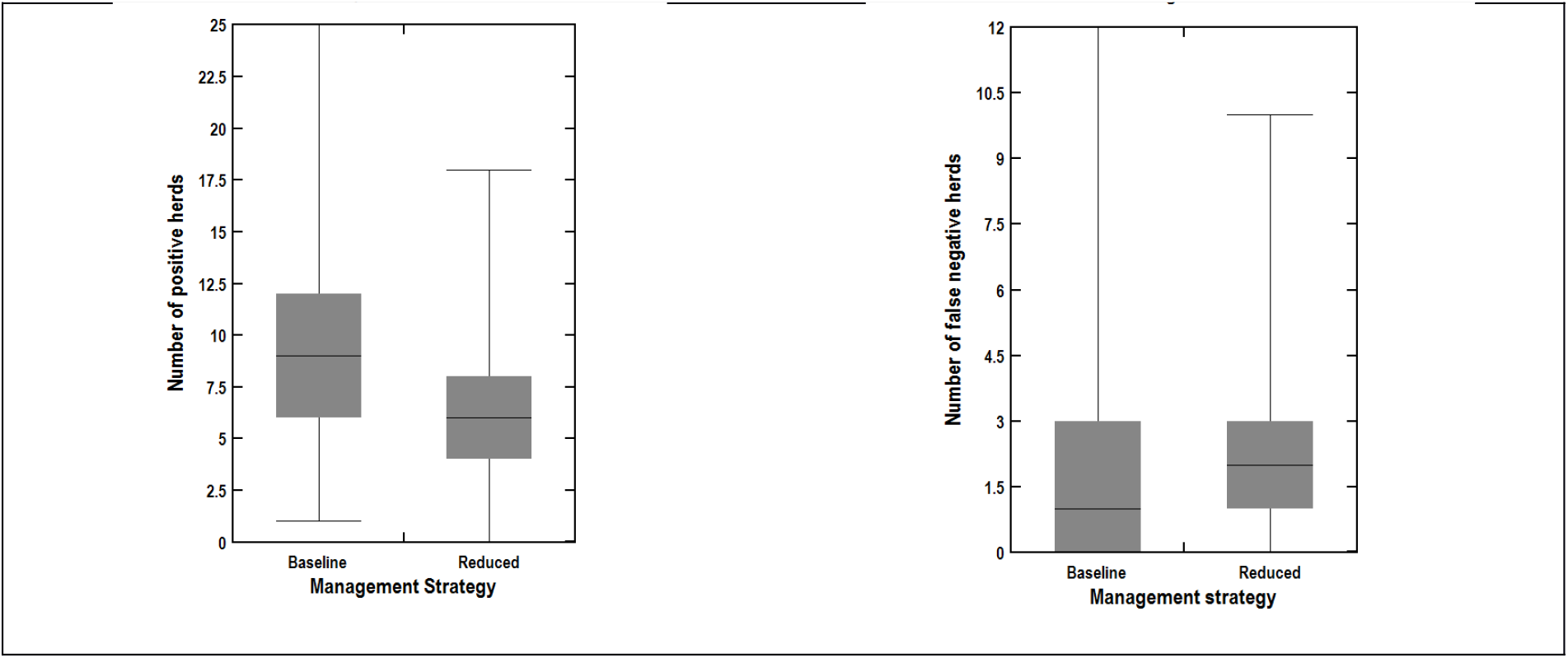
Comparison of baseline and alternate sampling regimes after vaccination control program

The reduced intensity sampling regime significantly reduced the number of herds tested, the number of samples collected, the cost of post-outbreak surveillance, and the number of positive herds requiring follow-up (on average by 7%, 49%, 24% and 50% respectively), compared to the baseline approach. The reduction in sampling intensity did not effect the number of false negative test results. Note that under both sampling regimes there is the possibility of small numbers of infected vaccinated herds being missed.

## 4. DISCUSSION

The focus of this study was on building modelling capability to quantify and compare the performance of different approaches to post-outbreak surveillance. This capability will support the development and refinement of policies on post-outbreak surveillance regimes to support regaining FMD-free status.

Limitations in sampling and diagnostic test performance mean that absolute freedom from infection is not provable in a target population (Schuppers et al., 2012). Instead, proof of freedom is substantiated through a surveillance regime that gives statistical confidence that disease is not present at a specified minimum level, or design prevalence (Paton et al., 2006). The probability of freedom in a herd can be estimated by sampling sufficient animals such that if no positive test results arise, a level of confidence is attained that the herd is not infected at the design prevalence. The design of a surveillance program is a compromise between the cost and logistics of implementing an intensive sampling regime, the cost-information ratio of additional samples, and the level of acceptable risk for the disease of concern (Schuppers et al., 2012). The level of acceptable risk may vary by country, by pathogen, and over time. For FMD, with its high economic impact, this is typically low, and is reflected in the design prevalence in protocols such as the EU FMD directive (European Union, 2003; Anon., 2007). It could be argued that in unvaccinated populations a higher design prevalence (with associated savings in time and sampling costs) would be adequate. For a highly contagious disease like FMD, it might be expected that a large proportion of an initially naive population would seroconvert to the disease, if it were present (Martin et al., 2007). The situation is less clear in vaccinated populations. The likely prevalence of infection in FMDV-infected vaccinated herds is not well quantified (Paton et al., 2006). Under some circumstances it could be quite low, so inevitably an intensive surveillance approach will be required.

Surveillance plans need to take into account the performance characteristics of the diagnostic tests. When testing a large number of samples, false positive results will be obtained, even using tests with high specificity and/or confirmatory tests. Simply removing reactors is not sufficient as their presence implies an overall failure to demonstrate freedom from infection (Paton et al., 2006). A surveillance plan should therefore include a follow-up protocol featuring resampling and testing for evidence of active infection with virological methods. Both the OIE code and EU Directive require that all herds with sero-reactors be followed up and classified as free or containing infection. Follow-up of reactors requires demonstrating the absence of transmission of FMD virus in vaccinated populations, where this is defined as ‘demonstrating changes in virological or serological evidence indicative of recent infection, even in the absence of clinical signs’ (OIE, 2016). The OIE code further states ‘in the absence of infection and transmission, findings of small numbers of seropositive animals do not warrant the declaration of a new outbreak and the follow-up investigations may be considered complete’ (OIE, 2016).

The case study demonstrated how the AADIS model can now be used to compare post-outbreak surveillance regimes, in previously infected areas, to support proof of freedom. The reduced sampling strategies were found to be more cost-effective than the baseline strategies. Compared to a baseline surveillance based on the European Union Directive, the reduced sampling intensity approach used after a control program, significantly reduced the number of samples collected and the cost of the post-outbreak sampling. There was also a significant reduction in the number of positive herds requiring follow-up. To recover FMD-free status, there should only be herds in the population that are seronegative or seropositive exclusively from the administration of inactivated vaccine. There should be no danger of carriers or undetected disease/infection. Under the assumptions used in this study, there were no residual herds under the non-vaccination control program, that is, the control measures put in place were effective in finding and removing all FMD-infected farms.

This was not the case when a vaccination-and-retain policy was used. With an emergency (suppressive ring) vaccination strategy, as applied in these studies, there is a high likelihood that some vaccinated herds will be exposed to infection before or soon after vaccination. Under these situations vaccination cannot be relied upon to prevent infection although it might suppress clinical signs in those herds. Residually infected vaccinated herds were not uncommon in the simulations. In the case study outbreak scenario there were up to seven infected and vaccinated herds present after completion of the control program. Post-outbreak surveillance programs cannot be guaranteed to find all of these herds and a similar number of herds were missed under both sampling regimes. Even with the baseline (EU) surveillance approach that involves testing all vaccinated animals, on average two true positive herds were missed in the case study outbreak scenario. ‘Small’ herds have been identified as a particular problem for FMD surveillance programs after emergency vaccination (Paton et al., 2006). This is because it is not possible to compensate for imperfect test sensitivity by increasing the number of animals tested. Options for dealing with small herds include (a) not vaccinating them in the first place (b) applying a vaccinate-and-remove policy for small herds (Paton et al., 2006; Anon., 2007). Animal health authorities should consider carefully whether vaccination of small herds is necessary under Australian conditions. In this study small herds were not vaccinated. It could be argued that small herds pose a relatively low risk of spreading infection, it is likely to be time consuming to vaccinate (numerous) small herds, and if vaccine is limited, larger herds would be a higher priority (Paton et al., 2006). Having said this, socio-political pressure could make it difficult to implement a non-vaccination policy for small herds under an emergency FMD vaccination program.

In this study, we have concentrated on structured surveillance in previously infected areas. This approach does not give an overall probability of freedom from disease. In practice a range of surveillance activities will contribute to the case for freedom from disease. A technique to take into account all the surveillance activities, such as scenario tree modelling (Martin et al., 2007), might help quantify the probability of being free from FMD.

## 5. ACKNOWLEDGEMENTS

This paper is derived from research undertaken by the authors in 2017 as part of CEBRA project 1604D ‘Incorporating economic components in Australia’s FMD modelling capability and evaluating post-outbreak management to support return to trade’. We would like to acknowledge the support of the Australian Department of Agriculture, Water and the Environment, the New Zealand Ministry of Primary Industries, and the Centre of Excellence for Biosecurity Risk Analysis within the University of Melbourne.

